# Mass Dynamics 2.0: An improved modular web-based platform for accelerated proteomics insight generation and decision making

**DOI:** 10.1101/2022.12.12.517480

**Authors:** Anna Quaglieri, Joseph Bloom, Aaron Triantafyllidis, Bradley Green, Mark R. Condina, Paula Burton Ngov, Giuseppe Infusini, Andrew I. Webb

## Abstract

Data processing is essential to reliably generate knowledge from proteomics studies. The complexity of the proteomics data, as well as the ability of research teams to adopt complex analysis pipelines, have proven to be an obstacle to effective collaboration and more efficient biological insight generation.

Here, we introduce MD 2.0, a cloud- and web-based platform for quantitative proteomics data, which implements a novel analysis workspace where statistical analyses, visualizations, and external knowledge generation are integrated into a modular framework. This modularity enables researchers the flexibility to test different hypotheses, and customize and template complex proteomics analyses, thereby expediting insight generation for complex datasets. The extensible MD 2.0 environment has been built with a scalable architecture to allow rapid development of future analysis modules and enhanced tools for remote collaboration, like experiment sharing and a live chat capability.

The new drag-and-drop modules allow researchers to easily and quickly assess different aspects of an experiment, including quality control, differential expression and enrichment analysis. The modularity of MD 2.0 lays the foundation to support broader community-based analytical template generation and optimized sharing and collaboration between proteomics experts and biologists, thereby accelerating research teams’ abilities to extract knowledge from complex proteomics datasets.

## INTRODUCTION

Advances in proteomics techniques, such as bottom-up liquid chromatography-mass spectrometry (LC-MS), have resulted in widespread adoption across the scientific community as the go-to tools to characterise complex biological samples. The versatility of LC-MS as an analytical technique has led to its uptake across fundamental research activities from basic science right through to drug discovery (including pre-clinical and clinical investigation), drug development (particularly in the biopharmaceutical industry), and in applied settings, such as food authentication, quality and safety^1–3^. The increase in its use has highlighted the need for developing reproducible sample preparation methods and more reliable hardware and software to ensure high levels of reproducibility and data integrity for quantitative proteomics analyses^4,5^. This is particularly important in MS-based clinical proteomics, where the robust profiling of various sample types, including tissues and biological fluids (such as plasma and urine) using LC-MS has demonstrated its potential as a mainstream clinical technique. The implementation of large-scale and possibly multi-institutional validation studies are now required to show the applicability of the chosen approach for use in clinical settings and yield protein assays with high diagnostic and prognostic accuracy^3^. However, these studies will require coupling with hardware for robust, fast, and automated sample preparation and data acquisition, as well as with software purpose-built to enable standardised processing of large datasets that allow multiple researchers with varying skill sets to accurately interpret the data produced.

To address this growing need, Mass Dynamics 1.0 (MD 1.0)^6^ was released in 2021 to provide a web-based data processing and analysis environment that is easy to use, reliable, and offers;

- On-the-fly, comprehensive quality control,
- Analyses of label-free quantitative (LFQ) proteomics data, and
- Straightforward collaboration and sharing capabilities.

The MD 1.0 platform allows the direct upload of pre-processed data from software such as MaxQuant, a Generic Format upload option for analysis of summarized protein intensities and on-demand processing for Thermo *.raw data-dependent acquisition (DDA). The original platform, MD 1.0, has been adopted across the globe since its introduction in 2019, with members spread across academic, government and industry settings.

Here we present the latest version of the platform, Mass Dynamics 2.0 (MD 2.0), which reframes the data analysis and insight generation process into a whole new modular framework that allows greater functionality through drag-and-drop analysis and visualisation modules and templates to create personalised workspaces. The modular and flexible framework was designed out of the necessity to support a diverse and growing field of researchers to: facilitate report personalisation, further standardise data processing, and increase the ease of integration of new data and analysis modules as they become available in the community.

MD 2.0 addresses and expands on some of the initial limitations of MD 1.0 around gene set enrichment analysis (GSEA) and collaboration capabilities. In particular, MD 2.0 extends the analysis and collaboration functionalities already available in MD 1.0 to support more bottom-up workflows thanks to:

- a more extensive visualization suite including interactive plots like heatmaps, upset plots, and a variety of linked plots such as volcano, violin, and dot plots;
- new data import options;
- the addition of gene set enrichment analysis (GSEA) with CAMERA^7^ via integration with the Gene Ontologies^8^ and Reactome external knowledge databases^9^;
- more direct collaboration between researchers by allowing note-taking in addition to sharing already available in MD 1.0.

The improvements in data import options in MD 2.0 allow the upload of data-independent analysis (DIA) files pre-processed with DIA-NN^10^ as well as the direct upload of DDA and DIA (via the integration of DIA-NN) Thermo raw files, extending the already available options in MD 1.0 for MaxQuant processed files or via a Generic Format upload. The new MD 2.0 platform can be accessed at app.massdynamics.com.

## METHODS

### Datasets

The LFQ DDA dataset with PRIDE identifier PXD016433 referred to here as “CKD dataset”^11^ (Chronic Kidney Disease dataset) was uploaded into the new interface via the Generic Format option. This was used to generate visuals and highlight knowledge integration. The dataset includes 36 separate samples extracted from the urine of patients with CKD. The samples belong to one of four conditions including CKD stage 1, stage 3 and stage 5, as well as control samples.

### DIA input workflows

Datasets acquired with DIA methods can be directly processed in MD 2.0 via integration of DIA-NN^10^. While we are continuously adding new capabilities, at the time of writing, data processing using DIA-NN is limited to library free searches and protein inference using protein isoforms. Data already pre-processed with DIA-NN can also be uploaded to MD 2.0 using the DIA-NN Tabular upload option. The outputs from both workflows are then post-processed to run quality control and statistical analyses and to allow the visualization of the results in MD 2.0. The downstream differential expression analysis is performed using the same workflow adopted for label-free quantitation data^12^.

### Mass Dynamics label-free quantitation (LFQ) data processing

The LFQ processing workflow uses the same approach as outlined previously for MD 1.0^12^ and summarised in the steps below:

- The raw intensity values are log-transformed with base 2;
- If Median normalization is applied, the median intensity of each sample is subtracted from the logged data;
- Missing values are imputed using MNAR imputation (1.8 standard deviations below the mean and with 0.3 standard deviation)^13^;
- Linear models are then fitted for each pairwise comparison using limma^14^. For each comparison, only proteins with at least 50% available measurements are considered.

### New interactive visualisations

MD 2.0 supports new interactive visualisations, including:

- **intensity heatmap**: a heatmap with hierarchical clustering of protein and samples intensities;
- **correlation heatmap**: a heatmap with hierarchical clustering of protein-protein correlations;
- **upset plot**: a type of visualisation used to describe the overlap between protein sets. The upset plot is generally preferred to Venn diagrams when more than three sets are considered;
- **dot** and **violin plots** which show the log2 intensities distribution of the selected proteins across the conditions in the experiment;
- **Reactome ORA strip and bar plots:** to comprehensively display the Reactome over-representation analysis (ORA) results. The strip plot is a dot plot displaying on the y-axis the -log10 (P-Value) for each gene set in the analysis while the bar plot shows the -log10 (P-Value) and the description of the 10 most significant gene sets.

The visualisations described above are generated in MD 2.0 through a service that reads the data uploaded, pre-processes it through one of available workflows, and generates visualisations upon request. The interactive plots are generated using Plotly (v5.3.0)^15^, Seaborn (v0.11.2)^16^, Matplotlib (v3.5.2)^17^ and UpSetPlot (v0.6.1)^18,19^.

### Enrichment analysis with external knowledge integration

MD 2.0 performs GSEA using the CAMERA method^7^. Camera is a competitive gene set testing method whose aim is to compare the differential expression ranking of the proteins (or genes) in a set against the ranking of proteins outside of the set, so as to highlight gene sets with highly ranked proteins. CAMERA is available as part of the Bioconductor package limma^20^ but in MD 2.0, it is accessed via the function, sbea (part of the R package EnrichmentBrowser), a wrapper for several available gene set testing methods^21^. In MD 2.0, gene set libraries are provided as input to sbea using the Gene Matrix Transposed (GMT) file format which contains the list of Uniprot IDs mapping to each one of the biological pathways tested. The GMT files are assembled from publicly available knowledge bases including UniProt, Gene Ontology (GO) and Reactome^8,22,23^. In particular, to assemble GMT files, the following data sources were used:

- UniProt: https://rtp.uniprot.org/pub/databases/uniprot/current_release/knowledgebase/complete/uniprot_sprot.dat.gz
- Reactome: https://reactome.org/download/current/ReactomePathways.txt https://reactome.org/download/current/UniProt2Reactome_All_Levels.txt https://reactome.org/download/current/ReactomePathwaysRelation.txt
- GO: http://purl.obolibrary.org/obo/go.obo http://current.geneontology.org/annotations/goa_human.gaf.gz http://ftp.ebi.ac.uk/pub/databases/GO/goa/MOUSE/goa_mouse.gaf.gz http://ftp.ebi.ac.uk/pub/databases/GO/goa/YEAST/goa_yeast.gaf.gz http://ftp.ebi.ac.uk/pub/databases/GO/goa/proteomes/264824.C_griseus.goa http://purl.obolibrary.org/obo/go.obo

Enrichment analysis in MD 2.0 is currently supported for the following species: Human (taxon id: 9606), Mouse (taxon id 10090), Chinese hamster (Cricetulus griseus) (taxon id 10029) and Yeast (Saccharomyces cerevisiae) (taxon id 559292). The source code for the enrichment workflow with example code and data is accessible at https://github.com/massdynamics/EnrichmentAnalysisStepR.

## RESULTS

With the introduction of MD 2.0, it is our aim to augment researcher capabilities to provide broader and richer access to the currently available analysis techniques as well as to create the foundation for a platform that can easily be extended with new functionalities as they are developed in the life science community. The major enhancements the enhancements over MD 1.0 are summarised in Figure 1.

**Figure 1:**
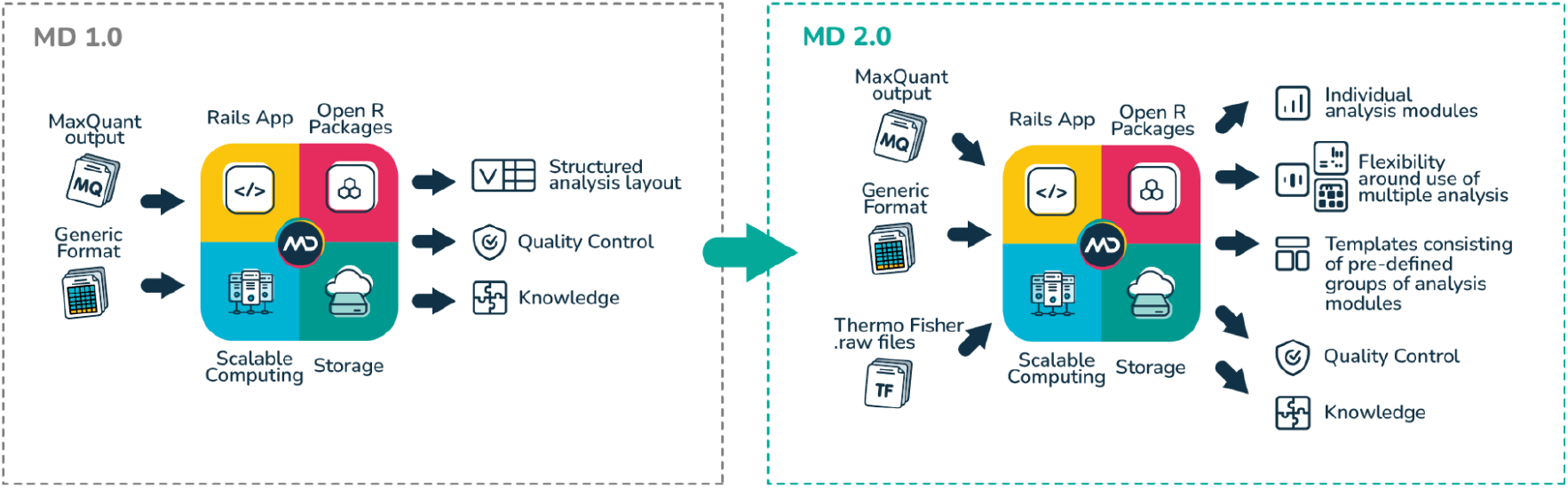
Overview of MD 2.0 functionality improvements including input options, user experience, data analysis capability and data management.

The “CKD dataset” (see Methods), was used as an example to highlight MD 2.0 functionalities in the sections below.

### Enhancements in MD 2.0: Increased User Functionality In Data Interrogation And Analysis

The new MD 2.0 platform takes inputs of experimental data in an expanded variety of file formats compared with MD1.0 (Figure 1). One major improvement in the MD 2.0 analysis environment is that the fixed analyses functionalities and visual representations of MD 1.0 are replaced in favour of a framework that allows researchers to explore the experiment results across various combinations of pre-defined analysis templates and modules to create customized workspaces.

Selecting templates automatically populates the tabbed workspace with pre-determined sets of analysis modules, which may be preferred to help speed up analysis and reporting, or those who prefer more rigidly defined experiment analysis flows. This gained flexibility unlocks the possibility of exploring a variety of hypotheses and generating multiple lists (of proteins or gene sets) that can be shared and assessed across multiple analyses and visualisations. In addition, the uncoupling of the analysis functionalities into separate modules is the building block that will allow easy integration with newly developed and user-requested modules.

#### Pre-defined Templates

In MD 2.0, researchers can modify and customize their tabbed workspaces and can delete or add modules. This new functionality allows researchers to customize modules to suit their needs and interrogate data in new ways. This approach is created by decoupling experiment data, protein lists and functionality. MD 2.0 has created specific templates that can be seamlessly generated, including templates for quality control, pairwise differential expression analysis, ORA, GSEA with CAMERA, heatmaps to show protein-protein correlation and sample’s clustering, and Upset plots to show intersections between protein sets (see Methods). The quality control templates display specific modules, depending on the input provided. A summary of the quality control template configurations are provided in Supplementary Table 1.

#### Modules

MD 2.0 introduces the concept of functional modules that relate to specific types of analysis that a user may wish to conduct (e.g., volcano plots, Reactome ORA etc). Further, researchers are now able to drag multiple analysis modules onto a workspace. In MD 2.0, tabs can be populated manually by selecting individual modules. These are grouped under functional categories and can be searched by name. A list with description of all the available modules is in Supplementary Table 2.

Using the new drag-and-drop functionality, researchers are able to drag modules and templates of interest onto a workspace contained in a tab. Additionally, they are able to create and name many workspaces (tabs), which allows them to effectively customize their experiment analysis by combining various analyses. Figure 2 shows an example analysis workspace in MD 2.0, where the tab is named “Heatmap and PCA” (Figure 2A), which was pre-populated to explore protein-protein interactions through a heatmap (Figure 2B), differential expression results and intensities distributions for selected proteins using the cross-linked volcano plot, results table and violin plots modules (Figure 2C-F) and clustering of samples using principal component analysis (PCA) (Figure 2G). Changes in one module, e.g. selection of the protein on the volcano plot, are reflected automatically in other modules via shared protein selections (and protein lists), e.g. the violin plots and results table are automatically updated. This cross-linking ability allows for faster interrogation of the selected proteins in the context of the whole experiment. Moreover, providing functionality in separate modules lays the foundation for researchers to create and make their own analysis algorithms available onto MD 2.0 for improved user experience and the added benefit of sharing it with the broader community.

**Figure 2:**
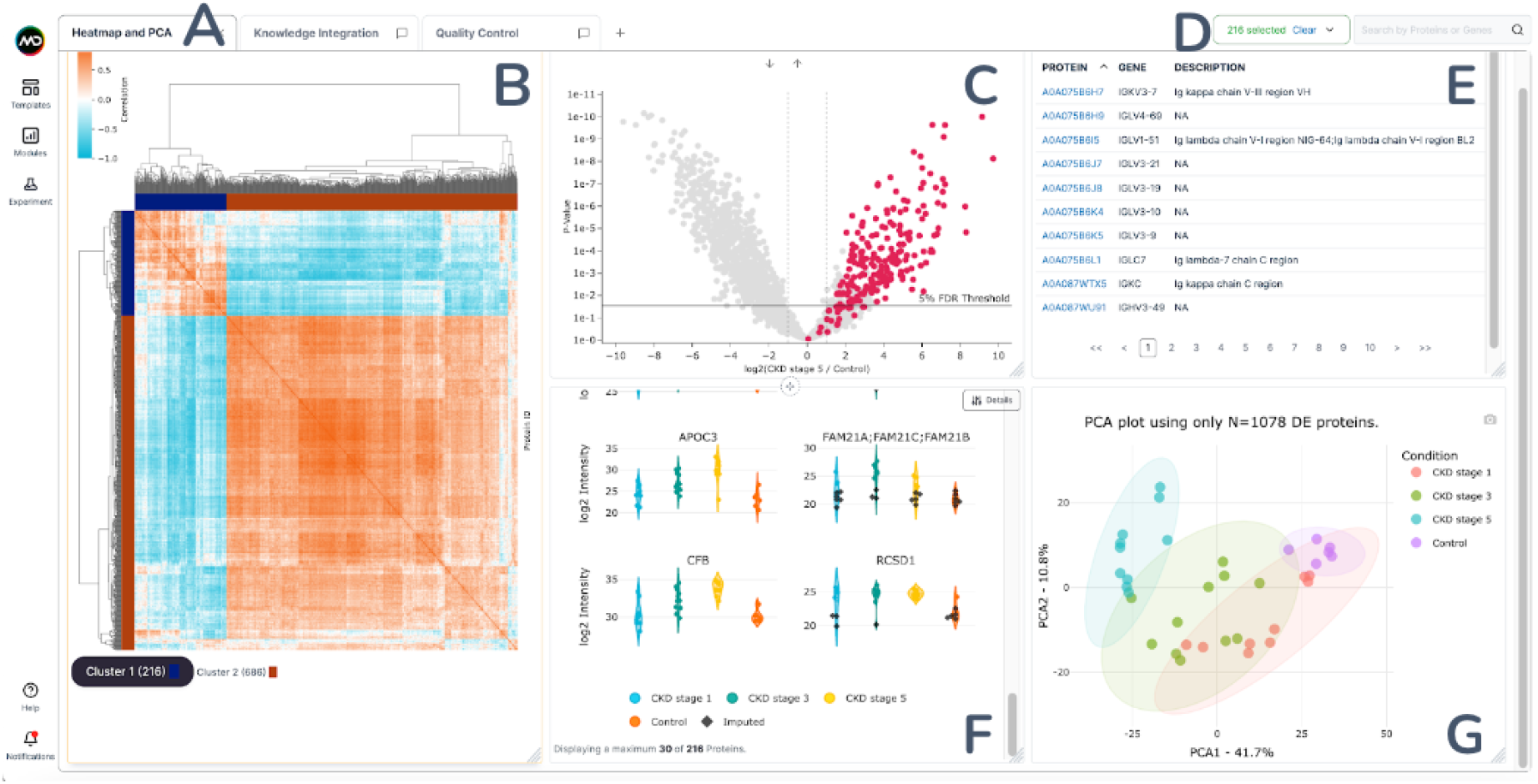
Improved user experience including new modules and templates for data interrogation and visualisation. (A.) MD 2.0 introduces the ability to create multiple workspaces (tabs) which provides the user with complete flexibility in separating their analysis activity groupings. (B-F.) MD 2.0 provides greater flexibility and customisation around analysis user flows and introduces ‘analysis modules’ which can be dragged into any workspace. (B.) Protein-protein correlation heatmap module. (C.) Volcano plot module. (D.) Data Table module of differential expression results. (E.) Violin plots module. (F.) Principal Component Analysis (PCA) module. (G.) Analysis modules can interrogate protein lists across multiple modules in real-time, when desired.

### MD 2.0 Improves User Experience with new Input Options

MD 2.0 has expanded the capacity of input options to support processing of DIA data. Researchers can now directly upload DIA-NN pre-processed data via the DIA-NN Tabular import option and access all the downstream statistical analysis and visualisations available in MD 2.0 modules and templates. Alternatively, raw DIA data can be directly uploaded in MD 2.0 and their processing is performed by leveraging DIA-NN^10^ prior to downstream statistical analysis. Furthermore, the MD 2.0 upload user experience has been improved for all import types by allowing the simple drag and drop of files and the intuitive assignment of files to different conditions so as to generate the experiment design. Future iterations will include in-app mapping of external data files, allowing researchers to upload almost any type of MS data file onto the platform.

### MD 2.0 Unlocks biological interpretations with External Knowledge Integration

A significant challenge in proteomic data analysis is interpreting results of a differential expression analysis which is often a list of proteins or peptides with large differences between conditions of interest. Often, this list is submitted manually to knowledge bases to interrogate potential relationships between proteins that have shown significant regulation between conditions. It can be difficult to define the relationships between proteins and how this relates to the experimental hypothesis.. MD 2.0 expands the interpretation capability by increasing the number of integrated external knowledge databases which now includes UniProt, Gene Ontology (GO) and Reactome^8,9,23,24^ and by adding a new service to test for the enrichment of significant biological pathways among the differential expression results. This allows researchers to interrogate observed proteins and how they relate to known biological categories (like GO and pathways) and determine observed proteins in an experiment that are related from a biological perspective, but are not differentially expressed.

#### MD 2.0 Integrates New Biological Pathways And Visualisations to the ORA service

MD 1.0 introduced the functionality for over-representation analysis (ORA) via the Reactome API content service, using protein lists generated from the pairwise comparisons results^12^. MD 2.0 now extends this service through a more comprehensive coverage of the Reactome database by including the disease and interactors databases and addition of new interactive visualisations.

Figure 3C shows a typical ORA analysis workflow in MD 2.0 using the “CKD dataset” (see Methods). In this example, a list of proteins is created using the clusters identified by hierarchical clustering through the heatmap module (“Cluster 2” highlighted in Figure 3A). Following this, the biological interpretation of the list is obtained with the direct interrogation of the Reactome database through the ORA modules. In particular, the significance of the top 10 gene sets returned by the ORA are displayed in a bar plot where bars are ordered by over-representation significance (Figure 3B) and the significance of all gene sets in the analysis are also displayed in a strip plot where the -log10 (P-value) is shown on the y-axis and the pathway considered on the x-axis (Figure 3C). The ORA results are also available as a table, which can be selected as another module not shown in the Figure (see Supplementary Table 2). Interestingly, this analysis reveals significant representation of pathways involving complement activation which was previously described in the “CKD dataset” original publication. This simple example shows how a few simple and intuitive steps involving straightforward protein selections and module interrogation can reproduce scientific findings which normally would require the ability to write several lines of scientific code and access to several scientific and visualization packages, i.e. heatmaps and extraction of clusters, ORA analysis, visualisation of results etc. Finally, each component of the ORA service can be used by itself as a separate module or they can be accessed together by selecting the Reactome ORA template.

**Figure 3:**
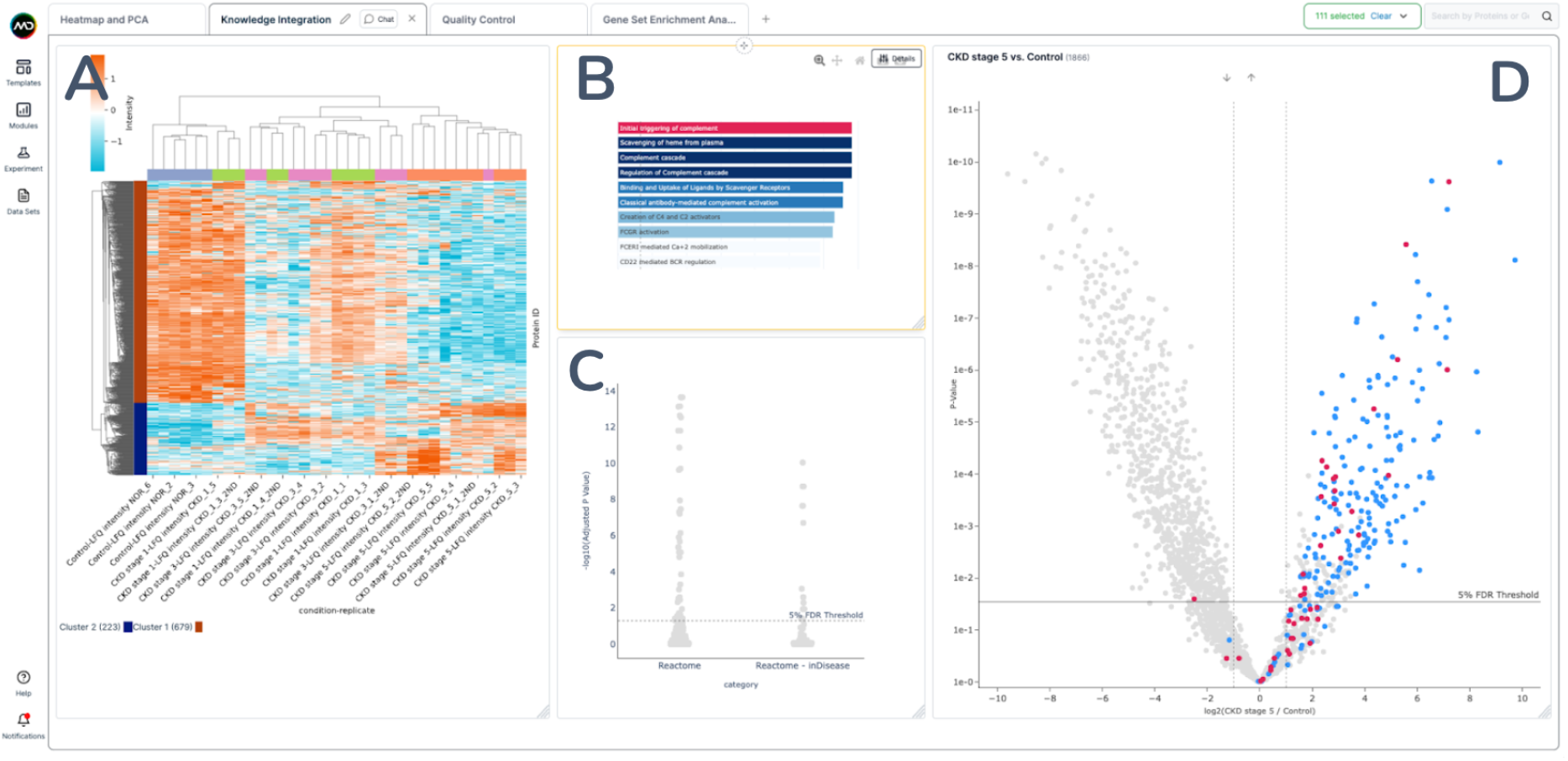
ORA analysis of PXD016433 - “CKD dataset”: Proteomic LFQ Analysis of Human Urine Samples (chronic kidney disease CKD stages 1, 3, and 5 vs healthy controls). LFQ data was uploaded via the generic format option and protein intensities were evaluated across samples using the generated heatmap. (A) The heatmap identified 2x main clusters. Cluster 2(223) consisted of proteins that sequentially increased with increasing CKD severity and as a result, this cluster was selected for ORA. (B) and (C) ORA analysis reveal significant representation of pathways such as complement activation, as previously described by Kim et. al^11^. (D) MD 2.0 allows researchers to link selected pathways seamlessly and overlay the proteins from the pathways with all associated proteins identified in the cluster in a pairwise comparison with controls.

#### MD 2.0 Enables Gene Set Enrichment Analysis With CAMERA

MD 2.0 integrates automated enrichment testing of Reactome and Gene Ontologies gene sets using the CAMERA method (see Methods) to enhance the interpretation of the differential expression (DE) results. While ORA simply looks for the over-representation of proteins from a particular biological set among a list of proteins provided (usually DE proteins) compared to all other proteins in the experiment, CAMERA takes into account the differential expression statistics obtained with the linear model framework, therefore testing for the up or down-regulation of each sets, and uses the estimated inter-gene correlations to obtain adjusted statistics. Figure 4 shows an example workflow analysis enabled by the MD 2.0 enrichment and cross-linking functionalities. In MD 2.0, the results of the enrichment analysis with CAMERA are gene set level statistics represented through linked results tables and volcano plots (Figure 4A-B). Each GSEA volcano plot (Figure 4A) is specific to a selected pairwise comparison and to a specific database, e.g. GO biological processes, and the gene set level statistics of all or selected gene sets can be searched and ordered using the GSEA results table (Figure 4B). The volcano plot shows, for each protein set, the -log10 (P-value) on the y-axis and Average Fold Change (AFC) on the x-axis. Selection of gene sets in the volcano plot also automatically filters the results table. Similar to the ORA functionality, researchers can generate protein lists directly from the results table by clicking the “View” option for a specific pathway and then “Select all proteins in gene set”. This action allows the creation of a new list which can then be interrogated across all other modules.

**Figure 4:**
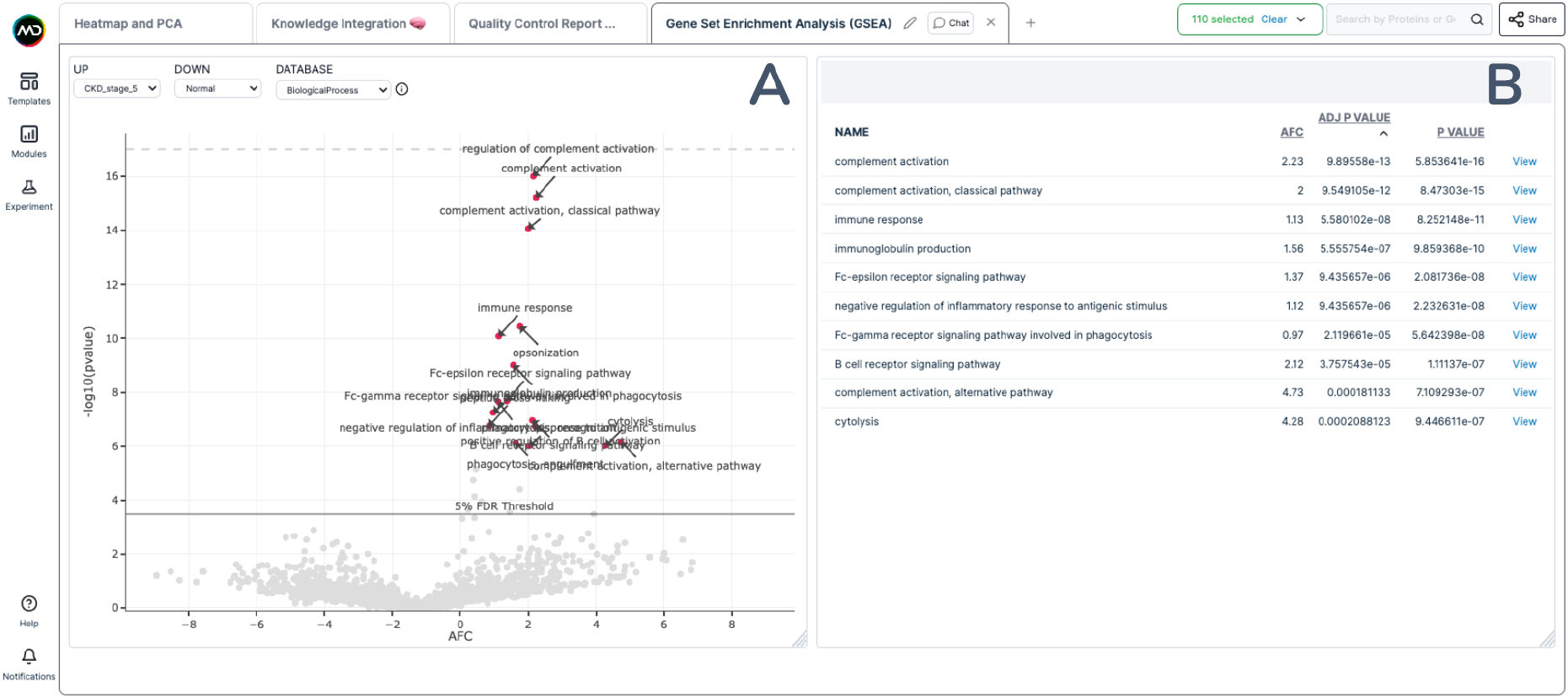
Example Enrichment Analysis Output using CAMERA with the PXD016433 data. (A.) GSEA Volcano plot showing the CAMERA results for the pairwise comparison “CKD stage 5” vs Normal and using the GO Biological Processes gene sets. The y-axis shows the -log10 (P-Value) and the x-axis the Average Fold Change (AFC) relative to each gene set. (B.) GSEA Results Table reporting the statistics for the gene sets selected in (A.) which includes: the name of the pathway, the AFC and the initial and adjusted P-Values.

### MD 2.0 Increases Data Management and Collaboration options

As per MD 1.0, experimental data is securely stored in the cloud and researchers are able to invite collaborators to work together on experiments and accelerate the gain of insights (Figure 5D). The security privileges provided to collaborators can be set to read-only access or full read-write permission. These functionalities are maintained in MD 2.0 but the modular layout lays the foundation for inviting collaborators access to only a specific tab workspace in an experiment. In addition to this, MD 2.0 elevates the collaboration options further with the integration of a workspace-specific chat box to allow real-time discussions directly in the experiment in app (Figure 5C). When a new message is sent, all collaborators with access to the experiment receive a notification by email and visible in the app (“Notifications” in bottom left corner of Figure 5) and are then able to reply directly using the chat. Furthermore, on top of the analysis modules, MD 2.0 provides text and checklist modules that enable note-taking and summary creation directly in the experiment workspace (Figure 5A-B, Supplementary Table 2). These new functionalities are designed to create a seamless link between data storage, analysis, results interpretation and insights sharing. This is to address the common bottleneck in multidisciplinary teams where discussions and information are usually spread and potentially lost across various files, emails and attachments, all of which tend to slow down interactions, insight generation and downstream decision-making.

**Figure 5:**
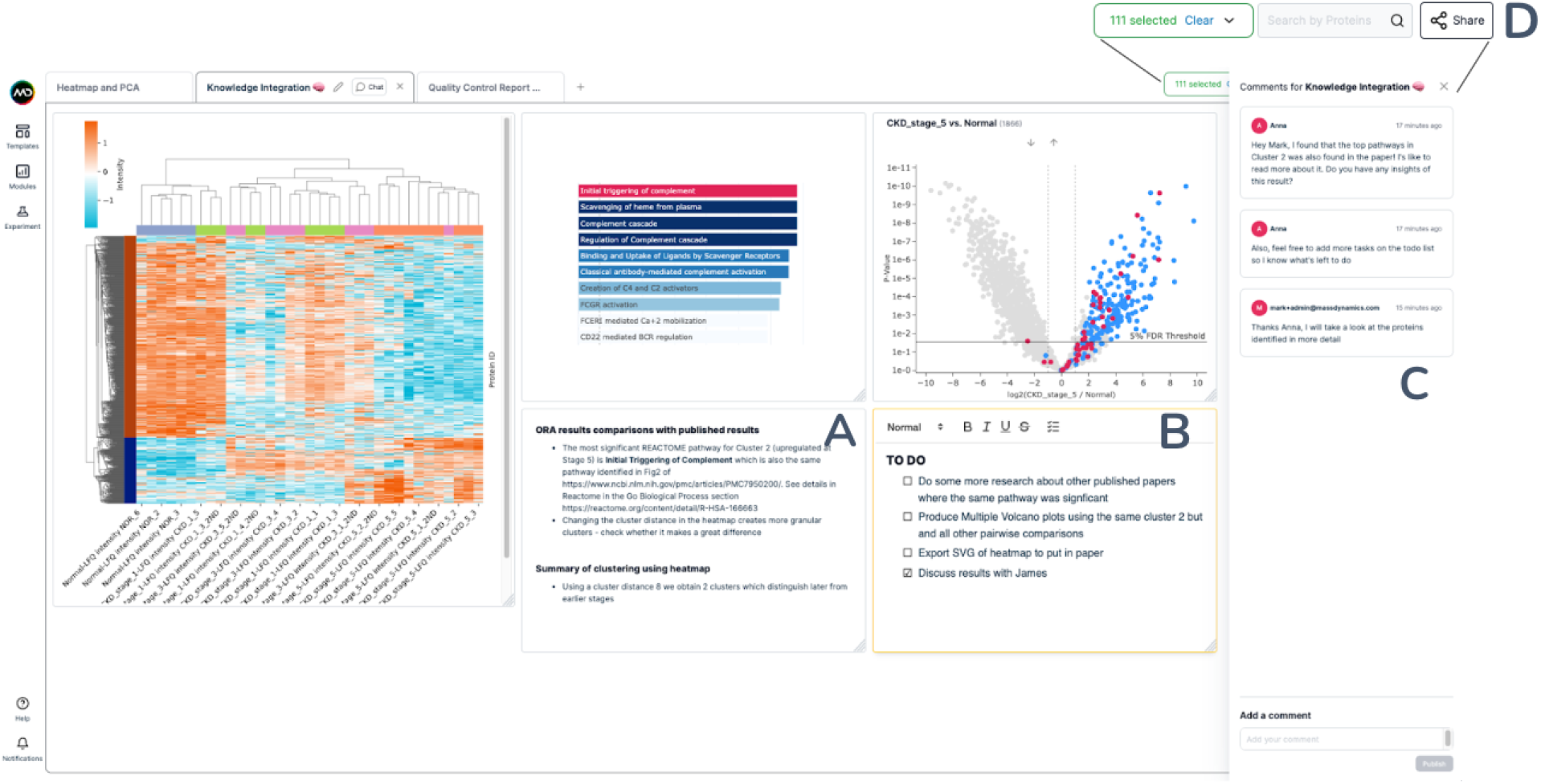
Example of enhanced management and collaboration options using the ORA analysis workspace of the PXD016433 data. (A) Text box to take notes. (B) Checklist box to create to-do lists. (C) Chat for live collaboration with collaborators that have access to the experiment. (D) “Share” button already available in MD 1.0 to share experiments with collaborators (covered by the live chat in the main panel).

### MD 2.0 Improves User Functionality through Real-time Advantage

Once processed and available in the app, the data from an experiment can be interrogated and visualised in the form of various visualisations (dot plot, list table, volcano plot, violin plot, log-log plots) to show the abundance and differential expression results of the quantified features as well as researchers can also access several types of analyses like ORA and GSEA with CAMERA. All of this occurs in real-time and with bi-directional feedback, enabling focused lists of proteins to be immediately interrogated across multiple analysis visualisations. As such, we use modern browser and framework capabilities, which unlock the capability of monitoring selected targets (e.g. proteins) in multiple visual analyses, as highlighted in Figure 2–5. This flexibility in user experience gives researchers both increased control and expanded data interrogation functionalities.

## DISCUSSIONS AND CONCLUSIONS

The recent rapid improvements in speed, reliability and depth of coverage of LC-MS based proteomics have led to significant growth in the volume and complexity of the biological measurements being generated. As a consequence, with the rapid expansion of proteomics data availability, it has become painfully clear that the most overlooked aspect of the analysis pipeline, generating insights, remains the most elusive and challenging as it is the most time-consuming component with the least access to automated tools. This part of the analysis has remained a multiple-step process, with the rigor of analysis depending on the level of understanding of the MS expert and/or biologist/s undertaking the work, as well as the types of bioinformatics workflows adopted to achieve the insights. In addition, new computational methodologies and workflows are continuously being developed to accommodate the needs of an expanding field (e.g. advancements in Single Cell and Top-Down Proteomics), making it necessary for anyone to be able to easily access and regularly integrate new data and analysis types.

While MD 1.0 has led the way with its web-based analysis platform to analyse, visualise and share LFQ data, this works demonstrates that MD 2.0, an enhanced modular web-based platform, simplifies, customises and templates complex proteomics analysis, facilitating rapid insight and knowledge generation from proteomics datasets. MD 2.0 has been conceived to offer a larger spread of scientific functionalities with human-centered design principles in mind, which is reflected in:

- The new flexible and customisable workspace,
- The new collaboration and upload options and
- The added functionalities for the biological interpretation of the results.

The web browser access for multiple collaborators coupled with the new flexible workspace experience and ability to interact in real time enables more efficient collaboration between proteomics experts and biologists, expediting decision-making. At the same time, the new modular framework lays the foundations for the integration of community-driven analysis templates so as to support the growing range of functionalities which are being developed. The new direct upload options for both RAW and pre-processed DDA and DIA data via an intuitive user interface allows simple drag-and-drop of files thus lowering the initial barrier of data preparation for upload and bringing the scientists closer to the relevant biological insights. Finally, the integration of an expanded set of external databases, including Reactome and Gene Ontologies, together with the addition of the GSEA with CAMERA, enables the seamless linking of LC-MS data to differential expression results and to their biological interpretation. The new scientific functionalities together with the ability to simultaneously interrogate different cross-linked modules and intuitively generate feature lists allows faster identification and evaluation of the mechanisms behind biological processes.

### Future development of MD 2.0

Although MD 2.0 addresses and builds upon the limitations of MD 1.0, future iterations are already in development for even more comprehensive user experience, functionality and analytic capability. Expansion of import pathways are in development to allow more seamless import from various sources. Dedicated templates for different experiment and analysis types are currently in development to give researchers greater flexibility and experience in using the MD 2.0 platform. The workflows supported for raw data in MD 2.0 is currently restricted to DIA and DDA label-free quant, while future iterations will broaden the scope of raw data that can be processed and expand on direct integration to external, publicly available, knowledge bases. Finally, label-based analysis (e.g. Tandem Mass Tags [TMT]) are currently only supported through the pre-processed options, such as with MaxQuant.

### Knowledge Generation is the Ultimate Goal

MD 2.0 accelerates knowledge generation and decision making by allowing faster and more intelligent interpretation of data and secure and seamless sharing of results to a multidisciplinary team, thereby informing validation studies and facilitating new hypothesis generation. In essence, MD 2.0 delivers a more powerful software platform with new functionality and an improved user experience. Leveraging MD 2.0 will help scientists elucidate the mechanisms of disease progression and facilitate the discovery of therapeutic targets.

## ACKNOWLEDGEMENTS

The authors would also like to thank the proteomics community for their ongoing development of comprehensive tools and repositories for the analysis and sharing of proteomics data because without them, the work in this manuscript would be impossible.

