## Supplementary Table 1 for "Mass Dynamics 2.0: An improved modular web-based platform for accelerated proteomics insight generation and decision making"

**Supplementary Table 2. Summary of MD 2.0 Improvements in Modules and Templates**

| **Name** | **Description** | **New or updated in MD 2.0** |
| --- | --- | --- |
| **MD 2.0 TEMPLATES can be predetermined or customised on request** | | |
| Quality Control Report | The QC Report template helps users assess several aspects of the experiment quality. It allows users to gain confidence about the health of their experiment prior to investing time in the analysis and results interpretation. | No change |
| Pairwise Analysis | The Pairwise Analysis template allows users to interactively explore the differential expression results by combining volcano and intensity distribution plots with the summary table. | Yes |
| Reactome over-representation analysis (ORA) | The Reactome ORA template combines several visualisations and a summary table to allow users to perform an Over Representation Analysis with user-selected protein lists to identify pathways relevant to the study. | Yes |
| Gene Set Enrichment Analysis (GSEA) | The GSEA template allows users to quickly run and explore enrichment results performed with the CAMERA method for several pairwise comparisons and knowledge databases. | No change |
| Heatmap | The Heatmap template combines Intensity and Protein correlation heatmaps. It allows users to quickly visualise protein expression patterns and cluster proteins that express in a similar way across samples. | Yes |
| Upset | The Upset template shows set intersections between user-created protein lists. | Yes |
| **MODULES:** Tabs can be populated manually by selecting individual modules. These are grouped under categories and can be searched by name. | | |
| **Experiment** | | |
| Experiment Design | A table listing groupings of Conditions and Filenames. | No change |
| **Pairwise analysis** | | |
| DE Summary | Table with the number of proteins included in each pairwise comparison after filtering for missing values and the number of DE proteins detected in the analysis. | No change |
| Volcano Plot | A plot to explore pairwise differential expression analysis. | No change |
| List Table | A table listing selected proteins, associated genes and descriptions. | No change |
| Data Table | A table showing differences between conditions including associated genes and descriptions, log2 ratios, p-values and adjusted p-values. | No change |
| Violin Plot | A plot to show the probability density distribution of the intensities of selected proteins, by conditions. | Yes |
| Dot Plot | A plot to show the distribution of the intensities of selected proteins, by conditions. | Yes |
| Pairwise Comparisons Log Log Plot | A 2D plot that shows the signed adjusted p-values on the log scale for two sets of selected pairwise comparisons, one comparison on the x-axis and one on the y-axis. The sign is derived from the log Ratio. | Yes |
| **Over-representation analysis (ORA)** | | |
| Reactome ORA Table | List of potentially over-represented pathways including estimated FDR and number of associated proteins. | No change |
| Reactome ORA strip plot | Scatter plot of potentially over-represented pathways ordered by -log10 estimated FDR. | Yes |
| Reactome ORA bar plot | Bar plot showing the top *n* potentially over-represented pathways ordered by -log10 estimated FDR. | Yes |
| **Gene set enrichment analysis** | | |
| Volcano Plot | A plot to explore the results of the pairwise GSEA, including the average fold change of the proteins in the set and the associated statistical confidence metrics. | No change |
| Results Table | A table listing the results of the selected gene set enrichment analysis. For each gene set, the corresponding absolute fold change (AFC), p-value and adjusted p-value are reported | No change |
| List Table | A table listing the selected enriched gene sets. |  |
| **Protein list** | | |
| Upset Plot | A plot showing the intersection of elements across user-defined protein lists. | Yes |
| **Quality control** | | |
| PCA | The Principal Component Analysis (PCA) plot is used to visualise differences between samples that are induced by the intensity profiles of quantified:  - proteins, modified peptides, peptides  - differentially expressed (DE) proteins, modified peptides,  peptides | No change |
| PCA Scree plot | A scree plot shows the amount of variance explained by each dimension extracted by the PCA of the quantified: - proteins  - modified peptides  - peptides | No change |
| CV Plot | Density plot of the % Coefficient of Variation (CV), obtained as the ratio of the standard deviation to the mean of the intensities for:  - proteins  - modified peptides  - peptides  and calculated by experimental condition. | No change |
| CV table | Table of median % CV by condition obtained for proteins (currently only available for the Generic Format workflow). | No change |
| Samples Correlation Matrix | Matrix that shows the intensity correlation between samples for:  - proteins  - modified peptides  - peptides  - DE proteins  - DE modified peptides  - DE peptides | No change |
| Sample Correlations scatter plots | Scatter plots that show the intensity correlation between samples for:  - proteins  - modified peptides  - peptides | No change |
| Feature completeness | Chart showing % missing values by:  - proteins  - modified peptides  - peptides | No change |
| Feature completeness by samples | Chart showing % missing values by samples for:  - proteins  - modified peptides  - peptides | No change |
| Missingness Patterns | Heatmap clustering samples by the pattern of missing intensity values (currently only available for the Generic Format workflow). | Yes |
| Identifications | Chart showing the number of identifications by sample for:  - proteins  - modified peptides  - peptides  - PSMs | No change |
| Consistent Identifications Proteins | Chart showing the number of proteins which are consistently identified by condition (currently only available for the Generic Format workflow). | Yes |
| Digestion efficiency | Barplot showing the % of missed cleavages observed.  Data obtained from the evidence.txt file after MaxQuant processing. | No change |
| Initial intensities boxplots | The boxplots of the log2 intensities before any normalisation or imputation is applied. Zero (missing) values are not included. | No change |
| Relative log expression (RLE) value plots before and after normalisation | The boxplots of relative log expression (RLE) values before and after normalisation (when applied). | No change |
| Imputation density plot | Density distribution of imputed vs actual intensities. | No change |
| **Heatmap** | | |
| Correlation Heatmap | Heatmap to assess protein-protein intensity correlations. | Yes |
| Intensity Heatmap | Heatmap that shows protein intensities across samples. | Yes |
| **General** | | |
| Text box | Write and format text to record insights and notes. | Yes |
| Checklist | Write, format and manage tasks. | Yes |
